# Reversible excision of the *wzy* locus in *Salmonella* Typhimurium may aid recovery following phage predation

**DOI:** 10.1101/2024.09.17.613263

**Authors:** Oliver JD Charity, Gaetan Thilliez, Haider Al-Khanaq, Luke Acton, Rafał Kolenda, Matt Bawn, Liljana Petrovska, Robert A Kingsley

**Affiliations:** Quadram Institute Bioscience, Norwich, UK; University of East Anglia, Norwich Research Park, Norwich, UK; School of Biotechnology, Dublin City University, D09 DX63, Dublin, Ireland; Department of Biochemistry and Molecular Biology, Faculty of Veterinary Medicine, Wrocław University of Environmental and Life Sciences, Poland; Earlham Institute, Norwich, UK; School of Natural and Environmental Sciences, Newcastle University, Newcastle, UK; Animal & Plant Health Agency (APHA), Weybridge, London, United Kingdom, UK

## Abstract

Bacteriophage (phage) are promising novel antimicrobials but a key challenge to their effective implementation is the rapid emergence of phage resistance. An improved understanding of phage-host interactions is therefore needed. The Anderson phage typing scheme differentiates closely related strains of *Salmonella enterica* serovar Typhimurium (*S*. Typhimurium) based on sensitivity to a panel of phage preparations. Switches in phage type are indicative of changes in phage sensitivity and inform on the dynamics of phage interaction with their host bacteria. We investigated the molecular basis of switches between the relatively phage sensitive *S*. Typhimurium DT8 and phage resistant DT30 strains that are present in the same phylogenetic clade. DT30 strains emerged from DT8 strains predominantly by deletion of a genomic region affecting the *wzy* locus encoding an O-antigen polymerase. The deletion site was flanked by two perfect direct repeats designated attL and attR. During broth culture in the presence of a typing phage that used O-antigen as primary receptor the Δ*wzy* genotype increased in frequency compared with culture in the absence of phage and removal of attL prevented deletion of the *wzy* locus. Co-culture of *S*. Typhimurium DT8 with a strain lacking *wzy* resulted in reversion of the latter to wild type. We propose a model in which reversible deletion of the *wzy* locus enables recovery of *S*. Typhimurium DT8 following predation by phage that use O-antigen as their primary receptor. This was consistent with ancestral state reconstruction of DT8 and DT30 phylogeny that supported a model of reversible transition from DT8 to DT30 in natural populations.

**Importance:** *S*. Typhimurium is a major pathogen of livestock that adversely affects productivity and animal welfare and poses a risk of foodborne disease in the human population. Antibiotics are used to control *Salmonella* infections in livestock that contributes to the antimicrobial resistance global emergency. Viruses of bacteria (phage) are one alternative to antibiotics to control *Salmonella* in the food chain but their successful implementation as antimicrobials is restricted by the rapid emergence of resistance to phage. A better understanding of the outcome of phage-bacteria interactions is needed to optimise the design and implementation of phage-based antimicrobials. This study identifies a genetic mechanism that confers resistance to phage that use O-antigen as a receptor on the surface of *Salmonella*. The mechanism is likely to impart a fitness cost on the bacterium but importantly the mechanism has the potential to be revert to a fully fit state once phage predation ceases. A model for how the mechanism may contribute to survival and recovery following phage predation is proposed.

## Introduction

The bacterial family Enterobacteriaceae include important bacterial pathogens of animals and humans including *Salmonella enterica* (1, 2) that are also a primary concern in the emergence of antimicrobial resistance (AMR). The extensive use of antibiotics in agriculture is thought to be a key factor in the emergence of AMR (3). Phages are viruses that specifically infect and kill bacteria and as such have potential as alternatives or adjuncts to antibiotics to treat infections, prevent transmission between hosts and reduce persistence in the environment (4–7) (8–11). A key barrier to the implementation of phages as antimicrobials is that bacteria that are initially sensitive to phage predation readily become resistant through spontaneous mutations affecting the expression of the bacterial cell-surface receptor used by phages to initiate infection. In our previous study, emergence of resistance within 18 hours of inoculation ranged from 10-100% of *S. enterica* serotype Typhimurium (*S*. Typhimurium) co-cultured with the single phage that were due to mutations that effected expression of lipopolysaccharide (LPS) on the cell surface (12).

Bacteria and phages exist in a continuous co-evolutionary arms race in which phage predation drives the evolution of antiphage systems (reviewed in (13, 14)) and phages evolve counter-measures to evade these systems (15–19). Phages require receptors which are constitutively present on the cell surface of bacteria for successful attachment and entry to continue the lytic cycle or undergo lysogeny. Sensitivity to phage predation is consequently heavily influenced by the presence or absence of these receptors (20). In laboratory experiments primarily investigating *Escherichia coli* interactions with various model phages, resistance rapidly emerged largely due to mutations affecting expression of the primary receptor (21). However, prolonged co-evolution has been observed, for example in batch cultures of *Pseudomonas fluorescens* involving multiple reciprocal selective sweeps of host and phage (22–24). Frequently phages target surface receptors such as lipopolysaccharide molecules (25, 26), whose mutation in many cases is accompanied by a high pleiotropic cost to fitness, limiting further co-evolution (27).

Evidence for the co-evolution and outcome of phage interactions with bacteria in nature is challenging and have therefore necessarily been largely limited to observational or correlational studies. This is in part due to the lack of large datasets linking phage sensitivity with bacterial genotypic diversity (24). A unique opportunity to investigate the dynamics of host genotype and phage sensitivity arose from the availability of a combination of phage type data and whole genome sequencing generated during surveillance of *S.* Typhimurium in the United Kingdom (28). Prior to the year 2014 the Anderson phage typing scheme was used to differentiate strains (phage types) of *S*. Typhimurium based on their sensitivity to 37 different phage preparations (29, 30). These phage types are stratified into more than 300 patterns of sensitivity which include 209 definitive types (DTs: DT1 - DT209), and over 118 undefined types (Us: U210 – U327) that have patterns not included in the original scheme. Whole genome sequencing (WGS) for surveillance of *S.* Typhimurium is now routine (31) and in the years 2014 and 2015 both phage typing and whole genome sequencing was performed on over 1700 *S.* Typhimurium isolates for surveillance and outbreak detection (28). The *<u>S</u>almonella* Typhimurium typing <u>p</u>hages (STMPs) group in three main clusters based on their sequence similarity. Most of the phage are P22-like phage and likely use LPS as primary receptor, STMP8 and 18 are ES18-like phage that likely target FhuA, while STMP12 and 13 are SETP3-like with an unknown receptor (32).

Phylogenomic analysis of *S*. Typhimurium sequence data revealed that major lineages were characterised by distinct patterns of sensitivity to phage indicated by the prevalence of specific phage types (28). An observation that was also true for enterohemorrhagic *E. coli* where phage type correlated with bacterial lineages (33). Furthermore, multiple lineages of *S*. Typhimurium containing multidrug resistant (MDR) strains that emerged and spread widely in livestock species over the past 50 years exhibited less sensitivity to predation by the panel of typing phages. There was also evidence for evolution to decreased sensitivity to the panel of phage during emergence and clonal expansion of the most recently emerged MDR monophasic (I,4,[5],12:i:-) clone of sequence type 34 (ST34) that emerged around 2005 in Europe (34, 35). The ST34 epidemic strains were predominantly DT193 in the year 2014 but ancestral state reconstruction indicated that the common ancestor of the ST34 epidemic clone was DT120 that had greater sensitivity to the panel of phage. Decreased sensitivity to the phage was associated with the multiple acquisitions of a prophage mTmII that resulted in clonal expansion of progeny perhaps due to the presence of an as yet unidentified antiphage mechanism (28).

Here we investigate the genetic basis of changes in sensitivity to phage predation in a sub-population of *S*. Typhimurium that are adapted to circulation in populations of duck and associated with occasional epidemics of human salmonellosis in the UK and Ireland (36, 37). *S*. Typhimurium isolates from this clade are predominantly DT8, with strains of DT30 that were characterised by a decrease in sensitivity to phage predation, sporadically distributed within the population structure of the clade. DT8 and DT30 are susceptible to STMP8, an ES18 like phage likely to target FhuA, an outer membrane protein. DT8 is also susceptible to STMP10, 11, 16, 20, 22, 23, 26 29 and 32, all of which are P22-like phages expected to target the O-antigen (32). We determine the molecular basis of the decreased sensitivity to phage predation leading to the change from DT8 to DT30. We test the hypothesis that precise excision of the *wzy* gene encoding an O-antigen polymerase is a mechanism of resistance and recovery following predation by phage that use O-antigen as receptor to initiate infection.

## Materials and Methods

### Bacterial isolates and whole genome sequencing

163 DT8 and 34 DT30 isolates were isolated during surveillance and obtained from the Animal and Plant Health protection Agency (APHA), and Public Health England (PHE) United Kingdom (now UKHSA). Whole genome sequence data for these isolates was produced by Illumina® Mi-Seq. Three DT8 clade long read reference genomes were (S04527-10 DT8, L01157-10 DT8, S03645-11 DT30) generated by Pacific Biosciences SMRT sequencing technology (38).

### Phylogenetic analysis

Phylogenetic trees from whole genome sequence data were constructed using SNPs identified in the whole genome sequences by aligning reads using BWA-MEM (39), variant calling with Freebayes (40) and SNP filtering using vcflib/vcftools (41), combined as a pipeline using Snippy v4.3.6 (42). Maximum-likelihood trees were constructed using a general time-reversible substitution model with gamma correction for amongst-site rate variation with RAxML v8.0.20 with 1000 bootstraps (43).

### Bacterial growth condition and recombinant strain construction

Bacterial strains were routinely grown in atmospheric conditions at 37□C on LB plate. Liquid culture were grown in the same condition with the addition of shaking at 200 revolutions per minute (RPM). Construction of recombinant strains was using the lambda red recombineering with primers GTGTAGGCTGGAGCTGCTTCG and CATATGAATATCCTCCTTAGT to amplify the *cat* gene or *aphII* gene from plasmids pKD13 or pKD14, respectively (44). Variable non-priming sequence at the 5’ end of primers was used to direct allelic replacement in the chromosome (sup Table 1). This approach was used to knock out the *wzy* gene (replaced by *aphII*), insertion of the *cat* gene into the *wzy-thrS* intergenic region, the attL*^wzy^* (replaced by *aphII* gene) region and to insert a kanamycin resistance cassette in the neutral intergenic region adjacent to the *iciA* gene in strain L01157-10 (45). The Lambda red recombinase encoding plasmid, pSIM18 was used for the recombineering step. This plasmid uses a temperature sensitive replicons and promotor to activate recombination machinery proteins *exo*, *beta,* and *gam* when exposed to 42□C, and cure the lambda red containing plasmids at 37°C. Knock in of a *wzy*-*tet* cassette, and *tet* cassette alone was carried out as described previously with minor modifications (46). A modified pDOC-GG-glmS plasmid with ampicillin selection marker was used as a delivery vector. The plasmid encodes a λ-red recombinase which allows sequence specific homologous recombination. The constructs were designed to insert the cassettes in the *glmS* intergenic region which was previously reported as a neutral locus for *S.* Typhimurium fitness (46). All the restriction enzyme digestion steps were using BsaI type II restriction enzyme (NEB). The Golden gate assembly of the modified pDOC-GG was using primers in Supplementary Table 1. The *tet* cassette was obtained by digestion of the muABbla tet 5’ plasmid. The cassettes were inserted in different *S.* Typhimurium strains as described in the text.

### Estimation of ancestral phenotype histories

We generated various models, and tested model fit by using the likelihood ratio test (LRT) in R. We compared scaled maximum likelihood generated ancestral probabilities at bifurcating nodes between models generated using an equal rate for transition (*Q*) or allowing different rates for *Q*. We also generated models using a Markov-Chain with Monte-Carlo (MCMC) sampling approach using equal rates for *Q* and another with a predefined distribution of *Qs.* The likelihood ratio distribution asymptotically converges toward a χ^2^ distribution. We could therefore determine which model was significant by using the LRT and identify where the result sits within a χ^2^ probability table, using one degree of freedom for an extra parameter of different rates of *Q.* The ancestral histories across a reconstructed phylogeny of 165 DT8 isolates and 34 DT30 isolates were assessed to determine the probable phenotype of each ancestor at each node in the tree. This was then used to determine a likely history of changes from the phage sensitive to phage resistant phenotype (DT8 to DT30). This was undertaken with maximum likelihood, and MCMC approaches (47). Maximum likelihood based states at nodes were estimated with Ancestral Character Estimation (ACE) from R package ape (48) and pastML (49). The transition rate matrices (Q) were estimated from the tip data and models allowing for different rates of transition were used. MCMC approach was conducted with discrete character mapping using posterior sampled maps from SIMMAP (50). One thousand sampled stochastic character maps were constructed after a burn-in period of 1000 iterations for *Q,* followed by 1,000,000 Markov-Chain steps and sampling for the posterior every 1000 generations using the pre-computed distribution of *Q* (51). Competing statistical models using either equal rates of transition or allowing different values for Q were compared using the likelihood ratio test (LRT, above) computed via an in-house script. The more complex model was required to reside in the right hand most 5% of a χ^2^ distribution with one degree of freedom to be considered significant. This was conducted using mean log-likelihoods of sampled estimated histories from MCMC analysis using both an equal rate Q, and a pre-computed values for Q estimated from the tip data. Resulting data was interpreted and viewed using R package phytools (51), iTOL (52), and pastML (49).

A permutation test was used to assess if the estimated character state at a node was due to a skewed result from sample bias, with 20 permutations. Tip labels were randomly assigned to the tree through the ‘sample’ R command and a starting tree constructed using the maximum likelihood-based function ACE software, with a subsequent tree sampled every 100 iterations of MCMC, 100 times, resulting in a distribution of 100 trees with ancestral states for each permutation. To assess whether the permutated data was from the same distribution as that estimated from actual tip data, pairwise, Mann-Whitney-Wilcoxon tests were performed.

### Bacterial genome wide association

Genome wide association was conducted to discern potential genetic polymorphisms associated with a trait of phage resistance for 34 DT30 isolates, or susceptibility to 10 phages for 164 DT8 isolates. This was undertaken through DNA of length K (k-mer) based analysis, where k-mers were identified from draft genome assemblies of each of the 196 isolate sequences using frequency based string mining algorithm FSM-lite (53). The approach adjusts the probability of association of k-mers based on phylogenetic structure. The population structure was estimated using mash and converted to a three dimensional distance matrix (54). Subsequently mixed linear model approaches were used for testing k-mer significance implemented with Sequence Element Enrichment (SEER) (55). This was done initially with no significance filtering of k-mers and then the top 1% of k-mers in the range of *p*-values determined a likelihood ratio test (LRT) *p*-value cutoff of 1×10^-3^. k-mers with a likelihood ratio test (LRT) *p*-value < 1×10^-1^ were plotted.

### Determination of sequence read mapping to the *wzy* locus depth

A genomic region of 6kb encompassing the *wzy* locus and flanking genes was extracted from reference sequence L01157-10 (RefSeq Accession no. GCF_902500305.1), and short-read Illlumina WGS data for 196 DT8 complex isolates was mapped to this region using Bowtie2 without filtering secondary mapped reads to ensure that any reads mapping to the region would be included (56). Bedtools-2.26.0 was used to extract the read depth per nucleotide (57). These were split into 250bp bins and the mean average of each section used as raw data for heatmap using R package gheatmap of ggtree (58). The read data was normalised by dividing the raw value for 250bp bins by the average read depth over the chromosome. This enabled visualisation of each *wzy* region in the context of each sequencing run. SNPs were identified in *wzy* regions using snippy v4.3.6 as previously mentioned with DT8 isolates S04527-10 and L01157-10 as references for variant calling (38).

### Determination of phage sensitivity and phage type

Phage typing was carried out as described in Public Health England’s phage typing protocol for *S*. Typhimurium using typing phages. STMP8, 10, 18, 20, 29, and 32 were a kind gift from UKHSA. Briefly, a single colony of the bacteria to be tested was incubated in 4 ml of nutrient broth with static, atmospheric conditions at 37°C for 2 hours. Subsequently a nutrient agar plate was flooded with the culture before drying. 10 μL of each phage suspension at recommended titre dilution was spotted onto the plate and incubated for 16 hours. Phage typing was used to assess that strains of DT8 and DT30 were susceptible/ resistant to the corresponding phage. Constructed mutants were also subject to challenge with the typing phage to assess the phenotype. A colony picked from the clearance zone in an experiment in which strain L01157-10 was exposed to phage STMP10 was used in reversion experiments (L01157-10 Δ*wzy*).

### Determination of genotype and *wzy* circularisation by PCR amplification

Presence of the *wzy* region was routinely tested using primer pair TGCGACTATCAGGTTACCGT and GTTAGCGTGCGGTCAAGATC that anneal in the *nucA* and *thrS* genes. qPCR was utilized to quantify the *wzy^-^*genotype in clonal populations with and without phage selection. Specific amplification of the circularised *wzy* locus was using primers AAGCCGAGACTCAGAGTGAC and CTCCGCCCTAATCCACATCT, and the same primers were used for the determination of the nucleotide sequence of the amplicon using TubeSeq (Eurofins).

### Quantification of *wzy* genotype

The relative and absolute quantification of *wzy* and Δ*wzy* genotypes was determined by qPCR using an Applied Biosystems™ StepOnePlus™. The strain to be tested was grown for 16-18 hours at 37°C in LB broth with shaking atmospheric conditions. This was sub-cultured in 10 mL of LB broth with 10^4^ cells per mL using a guide of an OD_600_ of 3.5 equalling 10^9^ cells per mL before challenge with 3.34×10^4^ or 3.34×10^5^ pfu/mL with STMP10. Then at 2-hour intervals for 8 hours, and a sample 24 hours post inoculum, 400 µL of culture was taken and stored at −20°C. Once all samples had been frozen genomes were extracted using Promega® Maxwell. L01157-10 was used as a positive control for growth, L01157-10 *wzy^-^* for control of growth of a resistant mutant under phage pressure, and a pure lysate with 3.44×10^5^ phage to control for any effect of phage DNA increasing the Ct values. qPCR was undertaken using extracted genomic DNA with *rpoD* as a housekeeping control gene, and *wzy* assessed using primer pairs wzy_qPCR_absent_F, wzy_qPCR_absent_R and wzy_qPCR_present_F, wzy_qPCR_present_R. A limit of detection for *wzy^-^* was established to the nearest log_10_ through 1 in 10 dilution of culture.

### Transfer of the *wzy* locus during co-culture

Co-culture and PCR amplification of the *wzy* locus from suspected revertant colonies was used to assess the reversion of *wzy.* A donor strain of *S*. Typhimurium L01157-10 *wzy^+^* was constructed by insertion of a *cat* gene conferring resistance to chloramphenicol into the intergenic region between *wzy* and attR^wzy^. A recipient strain of *S*. Typhimurium L01157-10 Δ*wzy* picked from the otherwise cleared plaque formed by phage STMP32 in an agar overlay assay and determined to have the deletion by analysis of whole genome sequence was further engineered by inserting an *aphII* gene conferring resistance to kanamycin in the 5’ intergenic region of *iciA* and subsequent selection of a nalidixic acid resistant variant by culture on LB agar containing 100 mg/l nalidixic acid. Two genetically unlinked selectable markers were used in the recipient to discount the possibility of transfer from the recipient strain to the donor. We reasoned that potential transfer of the *aphII* gene by transduction and selection for nalidixic acid resistant variants of L01157-10 *wzy^+^*are both rare events and therefore the frequency of both events occurring together is extremely unlikely. Presence of the *wzy* region was tested using primer pair TGCGACTATCAGGTTACCGT and GTTAGCGTGCGGTCAAGATC that anneal in the *nucA* and *thrS* genes, presence of *wzy* was tested using primers GCCTGAAGATTTTGGCGCAT, TGCGCTGACTTTGTTTCCTG that both anneal within the *wzy* gene, and presence of the *cat* cassette with ACAAACGGCATGATGAACCT and GCACAAGTTTTATCCGGCCT that both anneal within the *cat* gene. Six revertants were subject to WGS with Illumina HiSeq® 2500 to check the genotypes of the mutants and reversion of *wzy* within the same chromosomal location.

## Results

### The *S*. Typhimurium DT30 strains are sporadically distributed within the DT8 clade

Isolates typed as DT8 and DT30 are predominantly isolated from ducks and form a distinct clade within the *S*. Typhimurium population structure (28, 36, 38). Of 2213 *S*. Typhimurium strains isolated from ducks between 1992 and 2016, 68% (n=1509) were DT8 and 23% (n=352) were DT30. To further investigate the phylogenetic relationship of DT8 and DT30 strains we established a convenience collection of 176 whole genome sequences of *S*. Typhimurium isolated between the years 1993 and 2013 during surveillance by the Animal and Plant Health Agency (APHA), in the UK (Supplementary Table 1). These whole genome sequences were supplemented with 41 isolates from human infection and two genomes described previously (59). A maximum likelihood phylogenetic tree based on sequence variation in the core genome of 200 isolates (Figure 1A) indicated that strains of DT30 strains were distributed widely on the phylogenetic tree of the clade among DT8 with little evidence of clonal expansion.

**Figure 1.**
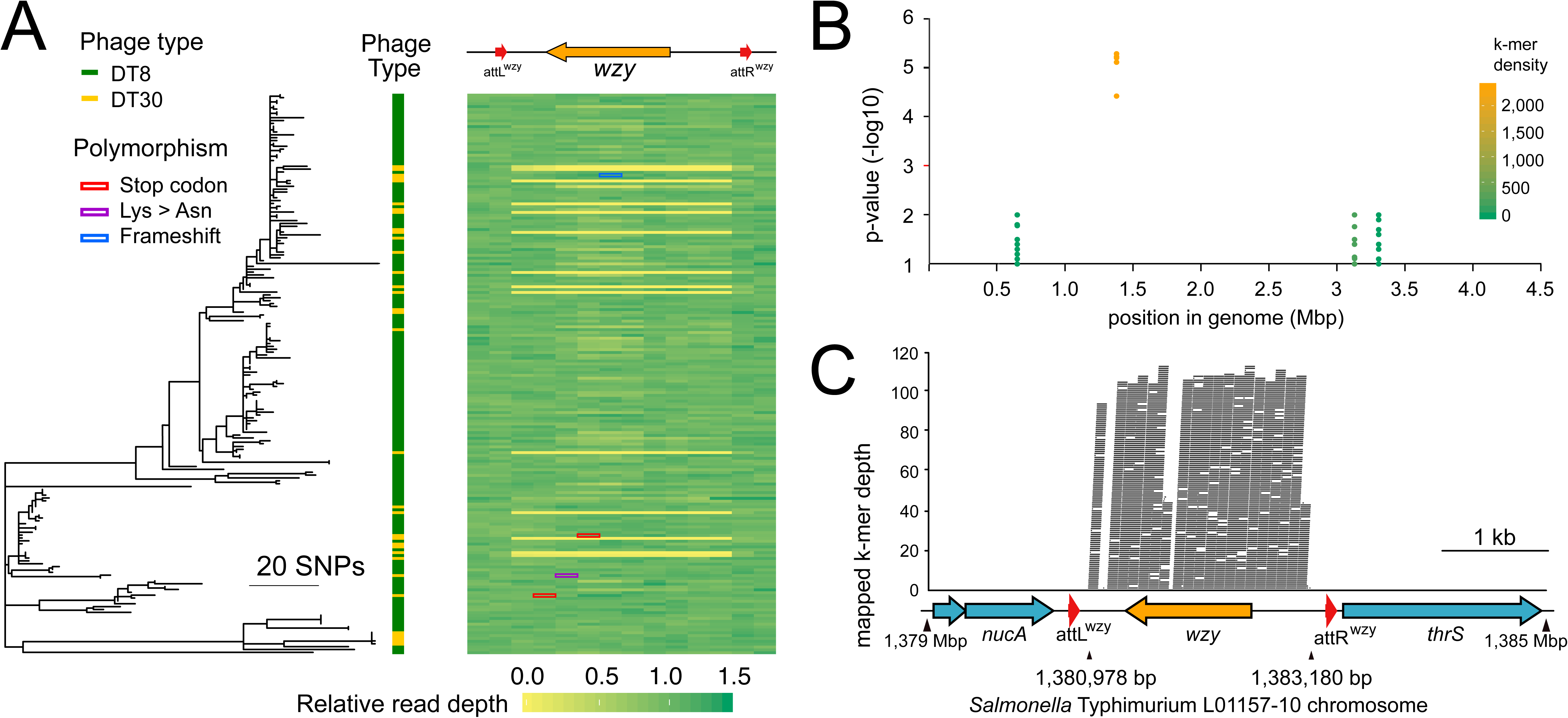
Phylogenetic relationship and association of sequence polymorphisms of *S*. Typhimurium DT8 and DT30 strains. (A) Maximum-likelihood phylogenetic tree constructed using 2,297 core genome variable sites from 162 DT8 strains and 34 DT30 strains. The heatmap indicates the relative level of reads mapped to the *wzy* locus of *S*. Typhimurium DT8 reference strain L01157-10 in 250bp bins for each strain. Coloured outlines denote the approximate location of polymorphisms that may disrupt Wzy function. (B) Bacterial genome wide association (GWAS) to identify sequence polymorphisms associated with phage resistance phenotypes of DT8 (susceptible) and DT30 (resistant). Manhattan plot of all significant k-mers position and likelihood ratio test (LRT) probability values of significant k-mers mapped to in *S*. Typhimurium DT8 reference strain L01157-10 chromosomal. The density of k-mers at a particular genomic coordinate is indicated by colour (inset key). (C) Location and depth of k-mers mapping to the *wzy* locus. Horizontal lines are k-mers with significant association with phage sensitivity defining DT8 and DT30

### The *wzy* locus is polymorphic in the DT8-DT30 clade and is flanked by two direct perfect repeats

To test the hypothesis that genome sequence polymorphisms account for decreased sensitivity to phage predation of DT30 compared to DT8 strains, a genome-wide association study (GWAS) was undertaken to identify k-mers associated with these two phagetypes. 2,617 k-mers that had a significant (likelihood ratio test, LRT p<0.1) association with the DT8 or DT30 trait after adjusting for the phylogenetic signal, mapped to four loci in the DT8 L01157-10 chromosome. A minority of k-mers (n=602) had an LRT p-value ranging from 0.1 and 0.01 and mapped to *yjbR* (n=200) encoding an uncharacterised DNA-binding protein encoding gene, *fucO* (n=201) encoding lactaldehyde reductase or *gss* (n=201) encoding a bifunctional glutathionylspermidine synthetase/amidase. The majority of k-mers, (n=2,015) mapped to *wzy* gene which encodes an O-antigen polymerase and had a likelihood ratio of association p-value < 0.0001 (Figure 1B). Quantification of whole genome sequence reads of the *S*. Typhimurium DT8 and DT30 strains that aligned to sequence within and flanking the *wzy* locus of *S*. Typhimurium DT8 strain L01157-10 whole genome sequence indicated a region of zero mapped reads in 14 of 34 DT30 isolates, consistent with a deletion of this region (Figure 1A). A further 20 strains that were typed as DT30 had sequence coverage across the *wzy* locus. Of these, three DT30 strains with sequence coverage had mutations resulting in a premature nonsense mutation in *wzy* and likely to abrogate function of the o-antigen polymerase, two SNPs and a small indel. Another strain had a non-synonymous SNP predicted to result in a lysine to asparagine substitution with unknown impact on function (Figure 1A).

The assembled sequence of *S*. Typhimurium DT8 strain L01157-10 has 2816 bp between the stop codon of *nucA* and the start codon of *thrS* compared to 259 bp in the assembled whole genome sequence of all 14 strains with the deletion at the *wzy* locus. Therefore, the deletion was 2558 bp. The genome sequence flanking the *wzy* gene contained two 113 bp identical direct repeats that coincided with the deletion boundaries observed in the DT30 strains in which the deletion at the *wzy* locus contained a single copy of the repeat sequence. The direct repeats flanking the *wzy* gene were therefore designated attL^wzy^ (*nucA* / *wzy* intergenic) and attR^wzy^ (*wzy* / *thrS* intergenic) and attB^wzy^ (Δ*wzy*) due to their resemblance to att repeat sequences that flank prophage in the bacterial chromosome and that mediate excision (60). Together these observations were consistent with a site-specific mechanism for deletion of this region (Figure 1C).

### Deletion of the *wzy* gene results in resistance to typing phage

To investigate the role of *wzy* in the sensitivity to predation by typing phages, a series of genetically engineered variants of DT8 strain L01157-10 in which the *wzy* gene was either deleted or replaced in an alternative genomic location for complementation were constructed. Strain L01157-10 Δ*wzy*::*aph*II in which *wzy* was deleted and replaced by the *aph* gene was resistant to phage STMP32 tested DT8-lysing phage (Figure 2). Reintroduction of the *wzy* gene into the 3’ intergenic region of the *glmS* gene, linked to a *tet* gene to aid selection, resulted in return of sensitivity to phage STMP32 while the *tet* gene alone did not result in restored sensitivity (Figure 2). The phage sensitivity phenotype of the L01157-10 Δ*wzy*::*aph* strain was therefore the same as DT30 strains. However, it is known that mutation other than those in *wzy* can result in changes in LPS expression that can also affect sensitivity to phage predation. We therefore tested if insertion of the *wzy* gene linked to a *tet* gene could also complement the phage sensitivity phenotype of strain S03645-11 DT30 that was resistant to typing phage STMP32 (Figure 2). Consistent with mutation of *wzy* in this strain being responsible for the loss of sensitivity to typing phage and the DT30 phenotype strain S03645-11 *wzy, tet* was sensitive to the typing phage. But introduction of *tet* alone resulted in no change to sensitivity (Figure 2). We therefore concluded that deletion of the *wzy* gene and flanking sequence was a common mechanism for the switch from DT8 to DT30 phage sensitivity phenotype.

**Figure 2.**
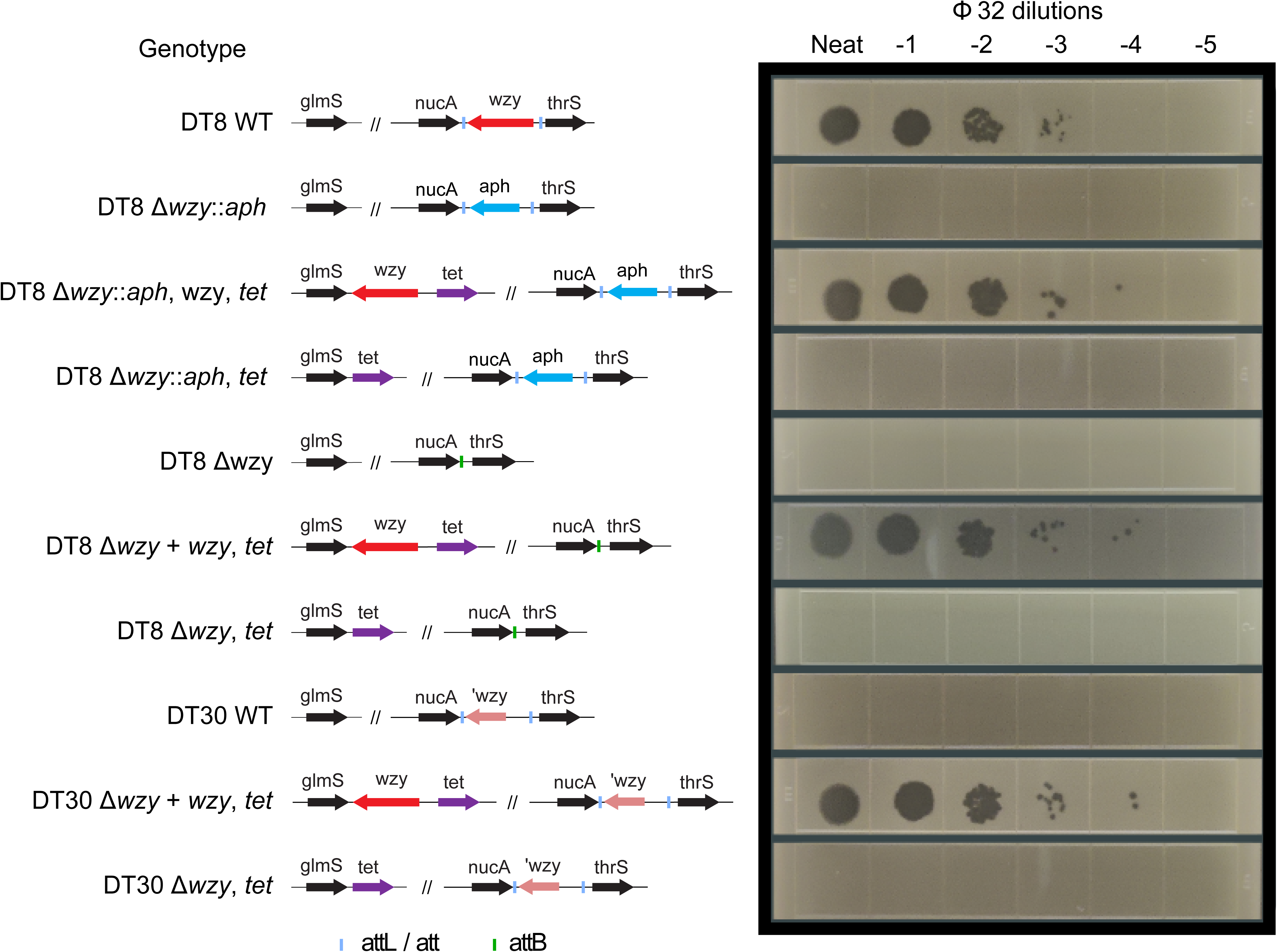
Sensitivity of *S*. Typhimurium DT8, DT30 and recombinant derivative strains to typing phage 32. The genotype of each strain is indicated with shaded arrows indicating genes as labelled (right) and images of serial dilutions of STMP32 overlaid on agar containing each strain.

### The Δ*wzy* genotype variants are present in cultures of *S*. Typhimurium DT8 that increase in frequency during phage predation

Since the *wzy* locus was flanked by attL^wzy^ and attR^wzy^ direct repeats was consistent with a site-specific mechanism of deletion we tested the hypothesis that excision of the locus occurs spontaneously during culture by amplifying by PCR across the *wzy* locus. PCR of genomic DNA prepared from stationary phase cultures of two DT8 strains L01157-10 and S04527-10 using oligonucleotide primers that flanked the *wzy* locus including the attL^wzy^ and attR^wzy^ repeats, resulted in two amplicons of 2.8 kb and 150 bp consistent with a mixture of the wild type *wzy* locus and the deletion variant. In contrast, a naturally occurring variant of L01157-10 in which the *wzy* locus had been deleted, picked from resistant bacteria in the zone of clearance due to lysis by phage STMP10 placed over a lawn of L01157-10 (L01157-10 Δ*wzy*), resulted in only the 150 bp amplicon consistent with deletion of the *wzy* locus (Figure 3A). Using qPCR with the same oligonucleotide primers we estimated that in stationary phase LB broth culture of L01157-10, the ratio of *wzy* : Δ*wzy* was approximately 500:1. To investigate the impact of phage predation on the ratio of *wzy* : Δ*wzy*, we determined the change in frequency of each genotype during culture of strain L01157-10 in the presence and absence of phage STMP10. In the absence of phage STMP10, the frequency of each genotype was stable during 24 hours of culture in LB (Figure 3B). In the presence of phage STMP10, the frequency of the wild type *wzy* locus decreased and the frequency of the Δ*wzy* genotype increased, consistent with selection of the deletion variant in response to phage predation. The genotype of a strain in which *wzy* was deleted and replaced by the *aph* gene remained constant in the presence of phage STMP10. Together these data suggest that the *wzy* locus is spontaneously excised during culture and the deletion variant is maintained at a constant frequency. Further, upon predation by a phage that uses LPS as a receptor, the frequency of the deletion variant increases relative to the wild type resulting in a change in ratio of *wzy* : Δ*wzy* to approximately 30:1.

**Figure 3.**
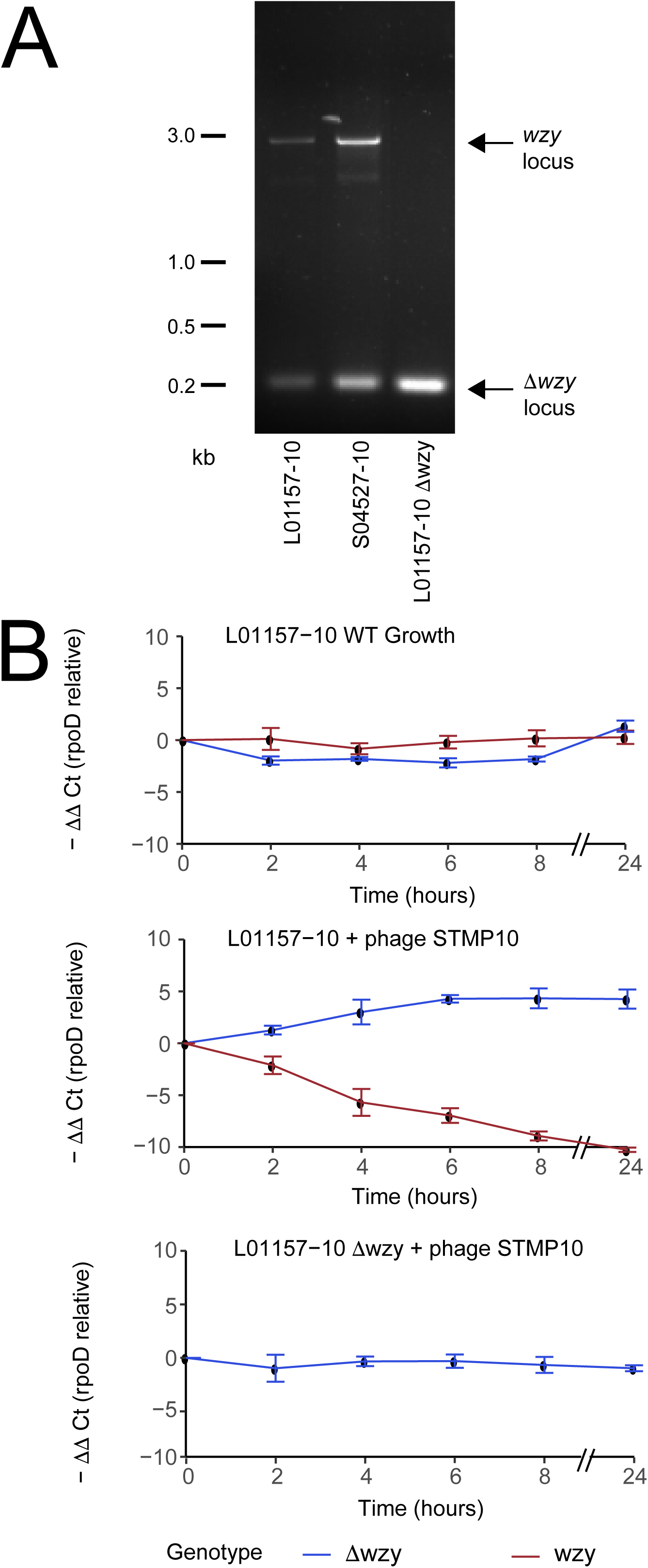
Determination of *wzy* locus genotype during culture and selection in the presence of phage predation. (A) PCR amplification of the *wzy* locus of two *S*. Typhimurium DT8 strains L01157-10 and S04527-10, and a spontaneous phage resistant variant of L01157-10. (B) Quantitative PCR of *wzy* genotypes of *S*. Typhimurium DT8 with and without pressure from STMP10.

### Deletion of the *wzy* locus is through an attL-dependent excision mechanism

To investigate whether the direct repeats attL^wzy^ and attR^wzy^ were required for excision of *wzy* locus, strain L01157-10 recombinants in which either attL^wzy^ or attR^wzy^ were replaced by the *aph*II gene were constructed by allelic replacement. An L01157-10 ΔattR^wzy^ strain had a growth defect during culture in LB as indicated by small colonies on LB agar, potentially due to the deletion affecting the *thrS* promotor region, and therefore was not investigated further. However, an L01157-10 ΔattL^wzy^ strain had a similar growth characteristic as the wild-type parent strain. PCR amplification across the *wzy* locus of strain L01157-10 resulted in a 2.8 kb band and a 150 bp amplicons but only the larger amplicon in L01157-10 ΔattL^wzy^ (Figure 4A). This was consistent with attL^wzy^ being essential for deletion of the *wzy* locus. Prophage excision that is also dependent on attL and attR repeats results in the formation of a circular molecular of the phage genome (60). To investigate if the *wzy* locus also formed a circular molecule, we PCR amplified genomic DNA prepared from strain L01157-10 using oligonucleotide primers that annealed within the excised region of the *wzy* locus in an outward facing orientation. This reaction resulted in a 0.5 kb fragment consistent with a circularisation of the *wzy* locus. The nucleotide sequence of the amplicon indicated that the amplicon spanned the left and right ends of the excised *wzy* locus consistent with a circular form of the *wzy* locus. Taken together, the observation suggested that loss of *wzy* is driven by excision of the region using the flanking attL repeat and likely the attR repeat (Figure 4B).

**Figure 4.**
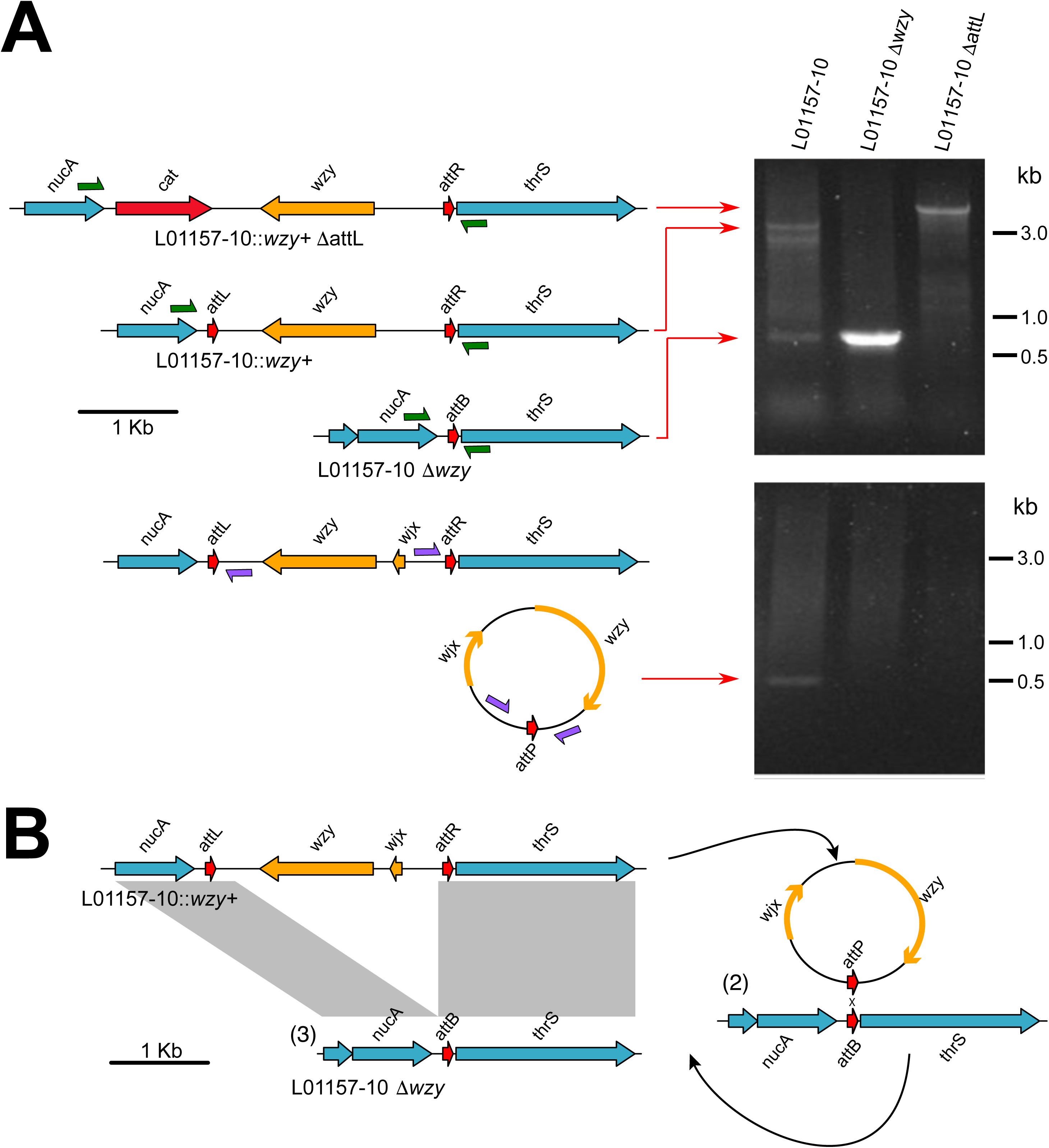
Dependency of *wzy* locus deletion and formation of a circularised *wzy* locus molecule on attL^wzy^ repeat sequence. (A) PCR amplification of the *wzy* locus from *S*. Typhimurium DT8 strain L01157-10, a spontaneous phage resistant variant of L01157-10 (L01157-10 Δ*wzy*) and a variant of L01157-10 in which the attL^wzy^ repeat sequence was deleted. Inward facing oligonucleotide primers (green arrows) flanking the *wzy* locus (top panel) and outward facing oligonucleotide primers (purple arrows) from within the deleted region (lower panel) were used in separate amplification reactions. Note: these are different primers than those used in Figure 3. (B) Model for the genotypes resulting from excision and circularisation of the *wzy* locus.

### Δ*wzy* deletion reversion to *wzy*^+^ during co-culture with a wild-type strain

We next investigated if Δ*wzy* can revert to wild type *wzy***^+^** during co-culture of a Δ*wzy* recipient strain with a donor strain encoding a wild type *wzy* gene. A donor L01157-10 strain was constructed in which the *cat* gene conferring resistance to chloramphenicol was inserted between the 5’ end of *wzy* and attR^wzy^ (Figure 5A, L01157-10 *wzy*, *cat*) so that transfer of the *wzy* locus could be selected by culture on chloramphenicol. A recipient L01157-10 Δ*wzy* strain that was spontaneously resistant to nalidixic acid and with an *aph* gene conferring resistance to kanamycin inserted in the 5’ region of the *iciA* gene by allelic exchange was used/constructed (L01157-10 Δ*wzy, aph,* nal^r^, Figure 5B). A mixture of donor and recipient strain cultured in LB for 24 hours resulted in chloramphenicol, kanamycin and nalidixic acid triple resistant L01157-10 at a frequency of 1×10^-9^ per CFU. Culture of the donor and recipient strain in LB supplemented with 0.5 μg/ml, that is known to induce prophage activation, resulted in an increase in frequency by 100-fold to 1×10^-7^ per CFU (Figure 5B).

**Figure 5.**
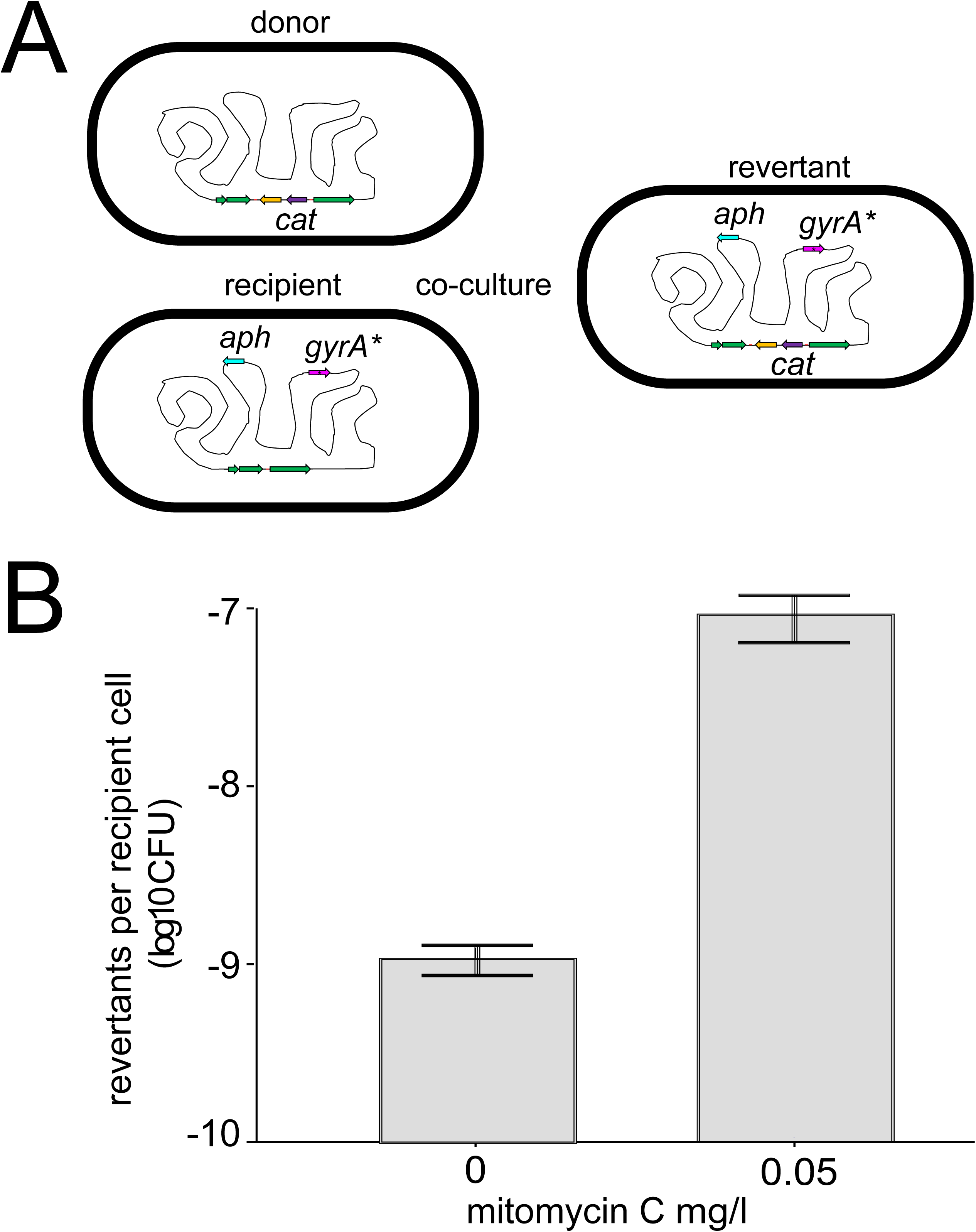
Transmission of *wzy* locus from *S*. Typhimurium wildtype L01157-10 to *S*. Typhimurium L01157-10 Δ*wzy* during co-culture. (A) Genotypes and genomic context of selectable markers and the *wzy* locus in L01157-10 *wzy*::*cat* donor, L01157-10 Δ*wzy aph*, Nal^r^ recipient and L01157-10 *wzy*::*cat aph*, Nal^r^ revertant strains. (B) Reversion frequency in the presence or absence of mitomycin C supplementation.

### Ancestral state reconstruction of DT8 / DT30 clade is consistent with bidirectional switching between states

Our laboratory data was consistent with a bidirectional excision and reversion of the *wzy* locus. To investigate the dynamics of phage sensitivity changes in natural populations of *S*. Typhimurium, we investigated the dynamics of DT8 and DT30 phenotype by estimating ancestral state at each bifurcating ancestral node and along each branch using two complementary methods, a Markov-Chain with Monte-Carlo sampling (MCMC) and a maximum likelihood approach. To account for varying rates of change in state we tested transition-rate matrices (*Q* matrices) to assess significance by testing model fit using the likelihood ratio test. For MCMC derived ancestral history probabilities, a *Q* allowing for different rates was significant over that with an equal rate *Q*, with χ^2^ with *p*=1.88×10^-15^. The model also had the greatest log likelihood of other models (scaled ML: equal rate *Q* = −143.6473, different rate *Q =* −90.9481, MCMC: equal rate *Q* = −140.3482, different rate *Q* = −185.748), and was therefore accepted as the most likely model out of all tested. To investigate whether the highest likelihood model where phage type transition rates were unequal was significant due to increased sampling of DT8 strains, rather than based on phylogenetic structure, permutation tests were undertaken. Twenty permutations were undertaken to assess if the probability distributions of each node being DT30 from each permutation were significantly different to the real data. A total of 18 permutations out of 20 were significantly different from the data (Supplementary Figure 1A). The Probability based ancestral estimation for the model with the greatest support indicated that ancestors were most likely to be DT8 (Figure 6A). A mean of 39.659 changes in phenotype state between DT8 and DT30 was estimated. The mean posterior values for *Q* were:

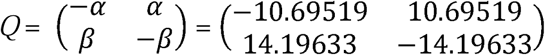

where

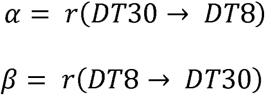

**Figure 6.**
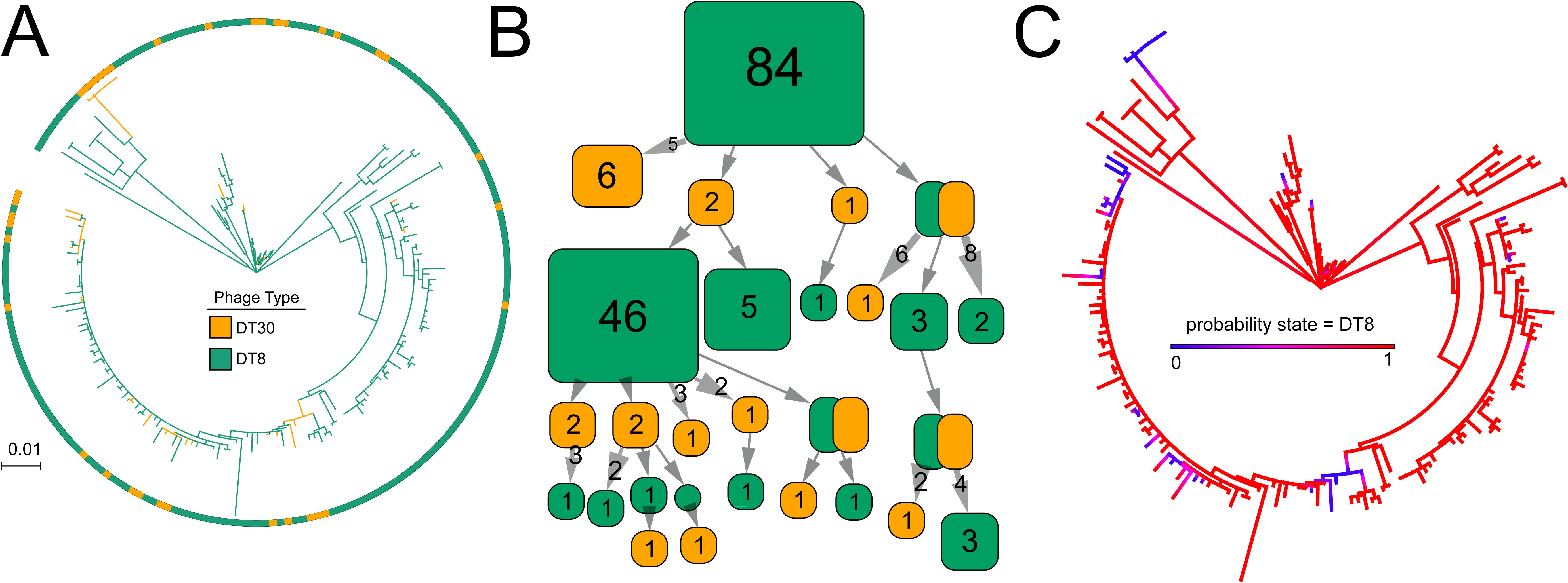
Ancestral state reconstruction of phage type for *S*. Typhimurium strains within the DT8 / DT30 clade. A) Maximum likelihood phylogenetic reconstruction of marginal ancestral states with empirical posterior probabilities used to infer an estimated history along the phylogeny. The outer ring indicated phage type DT8 (green) or DT30 (orange). The colours indicated at points in the phylogenetic tree are estimates of whether the state is bacteriophage resistant (DT30, orange) or bacteriophage susceptible (DT8, green). B) Collapsed-branch node graph to visualise the possible changes in state along lineages in (A). Filled squares indicate the state within a particular collapsed branch section of a tree, and how many tips track back to the section for DT8 (green) and DT30 (orange). Arrows indicate the direction of evolution, with the root node at the top and descendant nodes at intervals moving downwards. C) Probability density map indicating the probability that a state across the reconstructed phylogeny is DT8 from 1000 stochastic maps with periodic sampling from a distribution of possible trees with all DT8 probabilities shown as colours. Red indicates a probability of 1, and blue a probability of 0 that the ancestor was DT8.

These changes of state included an average of 25.754 switches from DT8 to DT30, and 13.905 for DT30 to DT8, which corresponds to the posterior values for rates of change between the two states.

In the complementary approach, maximum likelihood estimation of the probability distribution at each node indicated maximum marginal posterior probabilities similar that estimated using the MCMC approach (Figure 6B). A node graph indicated that most ancestral nodes were predicted to be DT8, but there were multiple predictions of transitions from DT8 to DT30 and back to DT8 (Figure 6C). A total of 45 changes in phage resistance phenotype were predicted, consisting of 26 switches from DT8 to DT30 and 19 switches from DT30 to DT8. The complementary approaches produced models that were highly correlated as indicated by an R^2^ of 0.675 for the probability of outcomes for each common ancestor being DT30 from scaled maximum likelihood ancestral states and stochastic mapping (Supplementary Figure 1B). Together these analyses are consistent with multiple state changes from DT8 to DT30, back to DT8 consistent with a reversable mechanism producing the resistant DT30 phenotype potentially occurring throughout the population.

## Discussion

Co-evolution of bacteria and phage is well-documented *in vitro* (22) and there is mounting evidence that phage predation impacts population structure (28, 33). Phage typing has been used historically to differentiate between closely related strains of *S*. Typhimurium and *S*. Enteritidis based on sensitivity to a panel of phage preparations (61). Investigation of phage type in the context of phylogenetic relationship of strains is one approach to understand the impact of phage predation on bacterial populations and the mechanisms that ensure fitness and survival. In a subclade of *S*. Typhimurium containing strains adapted to circulation in ducks, APHA surveillance data and our phylogenetic analysis of a subset of genomes indicated that around 80% were of DT8 and 20% DT30 that were sporadically distributed within the phylogenetic structure. The DT30 strains were characterised by a decrease in sensitivity to phage that use O-antigen as their primary receptor. In *S*. Typhimurium, less than 50-75% of isolates within epidemic clades have the dominant phage type (28) indicating frequent switching in phage sensitivity profile. In the case of the currently dominant pandemic multidrug resistant *S*. Typhimurium ST34 clone (35, 62), of 723 strains isolated from human infection in the UK in 2014, around 70% were of the dominant phage type DT193 (28). The remaining were of 14 different phage types. In this case the ancestor was predicted to be DT120, the second most common phage type in 2014. The switch from DT120 to DT193 was due to acquisition of a prophage, mTmII and resulted in a decrease in sensitivity phage and clonal expansion (28). Decreased sensitivity to phage predation may be a common characteristic of broad host range livestock associated *S*. Typhimurium, perhaps due to a fitness advantage from diminished phage predation (28). In contrast, the DT8/DT30 clade contained strains of DT30 distributed within the population of DT8, with little evidence of clonal expansion of the type with reduced phage sensitivity.

The reason for the difference in apparent fitness of DT30 investigated in this study and DT193 studied previously, as assessed by their relative level of clonal expansion (28) is likely to be the result the distinct mechanism of decreased sensitivity to phage. In this study, the *wzy* locus was deleted in 41% of DT30 strains, accounting for the change in sensitivity to phage using O-antigen as receptor and the change in phage type. Mutations in the coding sequence of *wzy* was also present in other DT30 strains. The *wzy* gene encodes an O-antigen polymerase required for the elaboration of long chain O-antigen on the surface that forms a barrier (63) that contributes to resistance to lysozyme (64), oxidative stress (65), colicins (66) and bile acids (67), and also affects the interaction with the complement system (68). It is therefore not surprising that mutation of *wzy* results in attenuation of virulence in mice, albeit not as attenuated as mutations in *waaG*, *waaI*, *waaJ*, *wbaP* and *waaL*, other genes involved in LPS biosynthesis that would also be expected to result in resistance to phage that use O-antigen as a receptor (69). In contrast, mTmII lysogeny responsible for the switch from DT120 to DT193 and decreased sensitivity to typing phage by an as yet unknown mechanism appears to have little or no fitness cost.

Genetic and phenotypic heterogeneity is a recurring theme in the interaction of bacteria with the host, environment and bacteriophage. The most common mechanism that generates heterogeneity in an otherwise clonal population is phase variation that results from slip strand mis-pairing, sequence inversion or differential methylation (reviewed in (70)). In bacterial pathogens heterogeneity generated from phase variation is important for functions including evasion of cross-immunity (71), persistence in the host (72), biofilm formation and detachment (73), bistable expression of energetically expensive type III secretion systems (74), and protection against bacteriophage and exogenous DNA (12, 75). The role of heterogeneity in defence against bacteriophage is particularly evident for genes involved in modification of LPS, a common receptor for bacteriophage. For example, it has been proposed that phase variation of the lic2A gene of *Haemophilus influenzae* encodes a glycosyltransferase that modifies LPS by the addition of galactose moiety is involved in a herd-resistance-like phenomenon resulting from reduced phage titres affecting predator-prey dynamics (76). *S.* Typhimurium also encodes multiple glycosyl transferase genes that modify LPS and an O-antigen chain length modulating gene whose expression are also under phase variation that contribute to resistance to bacteriophage (75, 77, 78). We describe a potentially novel form of phase variation in which heterogeneity is generated by reversible deletion of the *wzy* gene required to elaborate long chain O-antigen of LPS.

The mechanism for excision of the *wzy* locus is not known but we establish that it requires at least attL, one of a pair of 113 bp direct repeats flanking the *wzy* locus. The presence of attL and attR suggest that the mechanism may be analogous to other excision and integration systems that employ integrase and in some case with an excisionase (60). However, direct repeat sequences flanking the *wzy* locus are unlike the att repeat sequences of phage that are inverted repeats and often form attB and attC sequences with limited sequence similarity except for short (3-10bp) sequences where homologous recombination occurs. The *wzy* locus repeats bare some resemblance to the FRT and *loxP* sites of the site-specific recombinase systems from *Saccharomyces cerevisiae* and P1 phage, respectively, that can operate to excise or integrate sequence based on direct repeat sequences, catalysed by a site-specific recombinase. However, FRT and LoxP sequences are also distinct from attL and attR, each being only 34bp and composed of two symmetrical 13bp recognition sequences with a variable central sequence. In comparison, the *wzy* locus repeats are 113bp and lack symmetry within the sequence. We were unable to identify site-specific recombinase target sequences with a similar length and characteristics to that of the attL and attR. The relatively long and identical repeats may alternatively support the possibility that excision and integration was mediated by RecA dependent or independent recombination. RecA dependent recombination occurs between repeat sequences of greater than the minimal efficient processing segment (MEP) of 23-27bp (RecA/RecBC) and 44-90bp (RecA/RecF) with little increase in frequency above around 100bp (79). The greater the distance between repeats or decrease in sequence identity of the repeat sequence decreases the efficiency of recombination, and our observations are therefore consistent with such a mechanism. A RecA independent mechanism involving crossover between replicating sister strands, may also play a role (80), although it is not clear if this is capable of forming a circular excised molecular as observed for the *wzy* locus.

The potential for reversion of *S*. Typhimurium strains with the *wzy* gene deletion back to wild type by transfer of *wzy* from bystander *S*. Typhimurium is a critical observation that identifies this as mechanism that is potentially analogous to classical phase variation (70). The mechanism of *wzy* locus transfer is not known but the frequency increases in the presence of mitomycin C that is known to induce the SOS response and activate prophage (81). Prophage mediated genetic transfer has been observed in *S*. Typhimurium that was induced by the antibiotic carbadox (82) and prophage-like gene transfer agent (GTAs) in several bacterial species, that are thought to be cryptic prophage that retain the ability to package host DNA and carry out generalised transduction (83). Circularisation of the *wzy* locus in the donor strain may increase the frequency of transfer by protecting the molecule from endonucleases. Induction of the SOS response also increases expression of RecA and RecN that may be involved in homologous recombination at the attB site of the recipient and the attC site of the circularised *wzy* locus (84). However, it also remains possible that an integrase is responsible for this genetic event and this too has the potential to be up-regulated as part of the SOS response.

The observation that *wzy* deletion results in resistance to phage that use LPS as a receptor and the potential for subsequent reversion raises the possibility that this is a mechanism for recovery following phage predation. We determined that in populations of *S*. Typhimurium DT8 cultured in the absence of phage, approximately 1 in 500 had deleted the *wzy* locus, providing a pool of resistant variants that in the case of predation by phage using LPS as a receptor will survive the initial attack. However, *wzy* mutants have a fitness cost in the host and potentially in the environment and therefore reversion is key to this being an evolutionarily stable mechanism. To enable reversion, it is essential that *S*. Typhimurium with an intact *wzy* locus is also present following phage predation. Importantly, we determined that following culture with a phage that used LPS as a receptor, although the frequency of Δ*wzy* variants in the population increased to 1 in 30, still around 97% of surviving *S*. Typhimurium had an intact *wzy* locus. These were likely resistant to the phage due to mutations in other LPS biosynthesis genes such a single base insertion in the *pgm* gene that resulted in a frame shift predicted to truncate this gene (12). The potential for wzy^+^ and wzy^-^ variants to survive in the host or environment for long enough to allow for the predating phage to disperse is not known. During infections of the host, strains of *S*. Typhimurium with mutations in various LPS biosynthesis genes were able to colonise the Peyer’s patches, liver and spleen of mice following oral incoulation, but at lower levels compared to the wild-type strain. It is also noteworthy that mutation of *wzy* had the least impact of six genes that were tested on colonisation (69) suggesting that although Δ*wzy* variants may be present in only a relatively small proportion of the surviving population following phage predation, the *wzy*^+^ proportion may increase over time. The ability of strains harbouring mutations in various LPS biosynthesis genes to persist in the intestine is not known and this is an important factor that will determine the opportunity for reversion to occur. Another important factor is the dynamics of clearance of phage from the intestine once all phage sensitive strains have been killed and this is also currently not known.

Changes in O-antigen length due to phase variation of genes encoding chain length determination protein *opvAB* has also been described in *S*. Typhimurium and may represent a complementary system to that the *wzy* deletion (78). Phase variation results in a sub-population of *Salmonella* which have *opvAB*^on^ and *opvAB*^off^ allowing defence against certain phage that utilise LPS as a receptor, that was also recognised to come with an associated cost to virulence. In our study we did not investigate *opvAB* expression and therefore cannot compare the relative impact on evading phage predation with that of *wzy* deletion. It remains an open question as to how these two systems complement each other in evading phage predation.

The *wzy* deletion mechanism described is likely to be applicable to *Salmonella enterica* serovars of serogroups B1, D1 and A and at least some subspecies II serovars that encode an alternative *wzy* (*rfc*) encoded outside of the *rfb* locus (85, 86). Serogroup B1, D1 and A serovars are thought to have acquired the *wzy* in the *rfc* locus and the ancestral *wzy* in the *rfb* locus has been deleted leaving a sequence remnant (69, 86). Although only a subset of *S. enterica* serovars have the *wzy* (*rfc*) gene, these represent serotypes that account for the vast majority of human and livestock infections. For example, five of the top ten serovars accounting for non-typhoidal infections in people in the UK in 2022 are of serogroups B1 or D1 including Enteritidis, Typhimurium, Agona, Paratyphi B (Java) and Saintpaul, that together account for over half of all infections. Typhoidal *Salmonella* is a major cause of morbidity and mortality in South Asia, Africa and Latin America and *S*. Typhi and *S*. Paratyphi A that cause these infections both encode *wzy* in the *rfc* locus. Previous report of deletion of the wzy locus in strains of *S.* Enteritidis indicate that the deletion mechanism described in our study are likely to occur in other serovars with the *wzy* gene in the *rfc* locus (87).

In this study we investigated the deletion of *wzy*, the selection of a *wzy^-^* subpopulation during predation by phage that use LPS as a receptor and presented evidence of subsequent reversion to wild type. In this context we propose a working model in which phage predation results in the killing of sensitive *S*. Typhimurium and the survival of a subpopulation that have either deleted the *wzy* gene or have mutations in other LPS biosynthesis genes but retain the *wzy* locus (Figure 7). We propose that during infection or in the environment following phage predation, the absence of phage sensitive *S*. Typhimurium will result in clearance of the phage. The surviving subpopulation can therefore transfer the *wzy* locus from strains with alternate LPS mutations into Δ*wzy S*. Typhimurium reconstituting wild type *S*. Typhimurium. In this scenario, we would expect failure of phage treatment that could be tackled with repeated or continuous application of the phage. Alternatively, treatment with combinations of phage that exhibit collateral sensitivity due to the targeting of alternative receptors may also be effective in countering the proposed *wzy* deletion mechanism. These considerations are important to the effective deployment of phage as novel antimicrobials against *Salmonella*.

**Figure 7.**
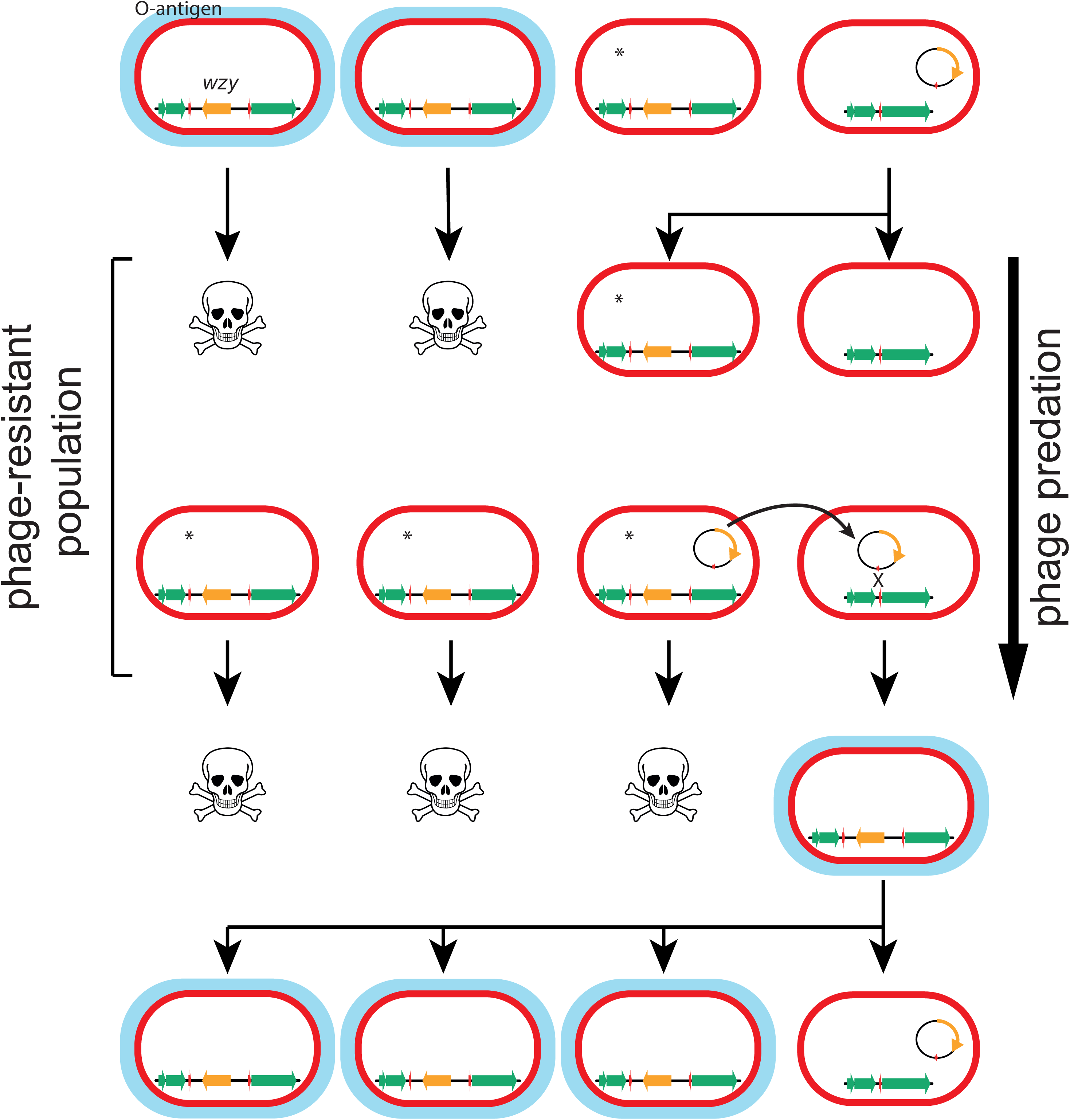
A model for how reversible deletion of the *wzy* may contribute to resistance to phage killing and recovery following transient predation by a lytic phage that recognises O-antigen as primary receptor.

## Conflict of interest statement

The authors declare that the research was conducted in the absence of any commercial or financial relationships that could be construed as a potential conflict of interest.

## Acknowledgement of funding

RAK laboratory was supported by research grants BB/N007964/1 and BB/M025489/1, and by the BBSRC Institute Strategic Programme Microbes in the Food Chain BB/R012504/1 and its constituent project(s) BBS/E/F/000PR10348 and BBS/E/F/000PR10349. OC was supported by a BBSRC DTP studentship (BB/M011216/1). LP was supported by Defra project RDOZO347.

## Notes

### Competing Interest Statement

The authors have declared no competing interest.

