## Supplementary figures and images for "Reversible excision of the *wzy* locus in *Salmonella* Typhimurium may aid recovery following phage predation"

### Supplementary Figure 1

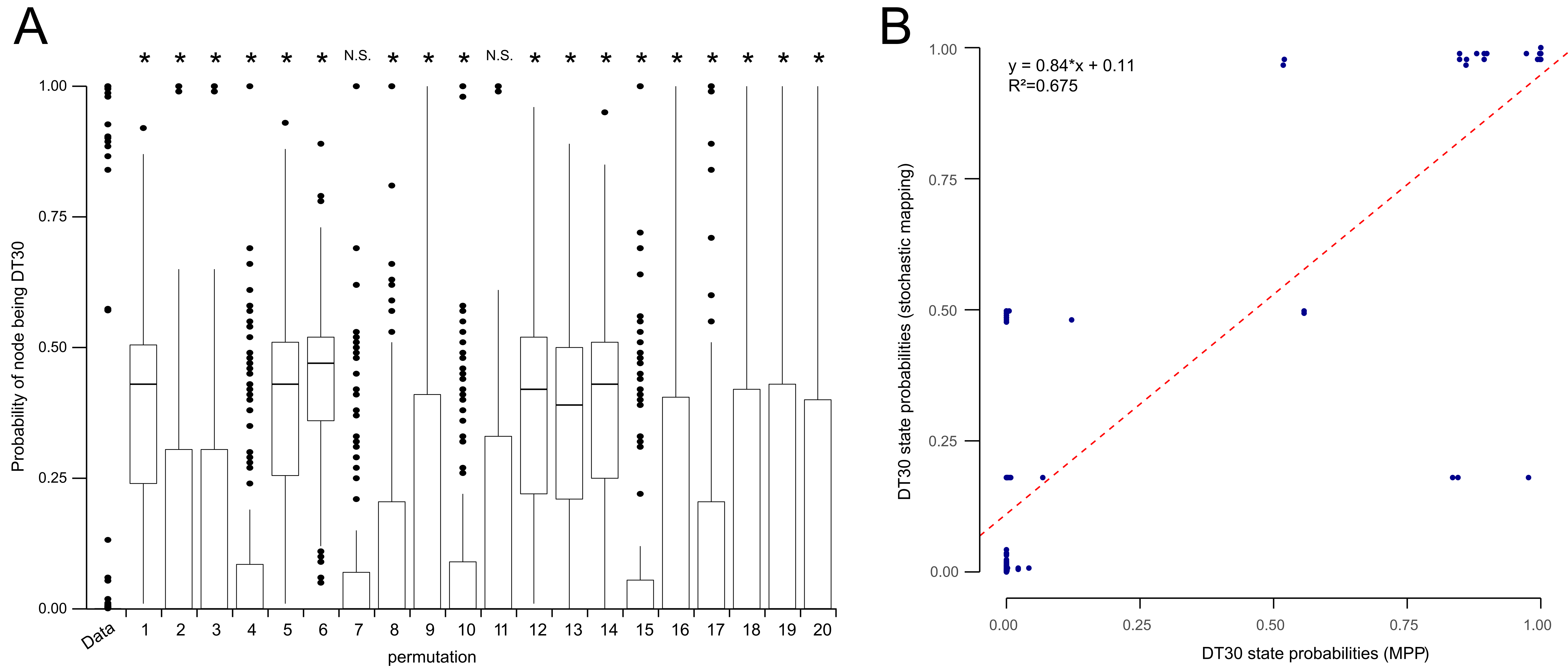
